# Characterization of a novel neurodevelopmental rare disease caused by a mutation within the autophagy gene *ATG9B*

**DOI:** 10.1101/2024.10.28.620020

**Authors:** Seval Kılıç, Kerem Esmen, A. Miray Oto, T. Bilge Kose, Melike Sever-Bahçekapılı, Emine Eren-Koçak, Şeyda Demir, A. Semra Hız, Gökhan Karakülah, H. Alper Bağrıyanık, Mehmet Öztürk, M. Kasım Diril

## Abstract

Autophagy is a highly conserved eukaryotic cellular process whose dysfunction results in human pathologies including cancer and neurodegenerative disease. First identified in yeast, ATG genes are central players in autophagy. Although their roles in cancer and neurodegenerative disease are well known, Mendelian diseases associated with ATG genes are rare. Mutations in core autophagy genes *ATG5* and *ATG7* have been previously reported to cause rare genetic disorders with autosomal recessive inheritance pattern.

Here we report, for the first time, a rare genetic disorder that results from a deletion/frameshift mutation in human ATG9B, the placenta specific homologue of yeast ATG9 in humans. The 11-nucleotide deletion causes a frameshift and addition of a premature stop codon, truncating the C-terminal cytosolic domain of the ATG9B protein. The pediatric patients carrying the mutant allele homozygous were children of a consanguineous marriage and displayed neurodevelopmental anomalies including mental retardation. We hypothesized that this phenotype originates during placental development.

To characterize the effects of the mutation and gain insight on the specific functions of ATG9B in a physiological setting, we used mammalian cells and generated a knock-in mouse model. Truncated ATG9B was not stable when expressed in cells. It was localized to perinuclear vesicles like the WT protein, but not to peripheral vesicles. Homozygous knock-in mice were viable, fertile and displayed no gross phenotypical abnormalities. Histomorphometry analysis of the placenta layers did not reveal a significant difference between mutant and control embryos. The assessments of neurobehavioral tests were similar in wild-type and homozygous knock-in mice. However, knock-in mice had a reduced fear memory trend, which is an amygdala-involved response.

## Introduction

Autophagy is a conserved catabolic process in eukaryotes, essential for intracellular degradation and homeostasis. Misfolded proteins, aggregates, and dysfunctional organelles are encapsulated by a double membrane structure forming autophagosomes and delivered to lysosomes for degradation (Yang and Klionsky 2020). The resulting monomers, such as amino acids, are then released into the cytoplasm for metabolic reuse or biosynthesis. Autophagy plays critical roles in morphogenesis, aging, cell differentiation and death, and intracellular immunity by degrading pathogens. Dysregulation of autophagy has been linked to pathologies such as cancer and neurodegenerative diseases, including Huntington’s and Parkinson’s disease (Cheng et al. 2013). Macroautophagy (commonly referred to simply as “autophagy”) involves the formation of autophagosomes that sequester large cargoes, such as protein aggregates and damaged organelles, which are then fused with lysosomes for degradation (Oral et al. 2017).

Autophagy process involves approximately 20 core autophagy-related proteins (ATG) mediating complex membrane dynamics. Yeast ATG9 and its mammalian homologs ATG9A and ATG9B are the only transmembrane proteins involved in autophagy (Matoba and Noda 2021). Autophagosomes formation begins by *de novo* fusion of Atg9 vesicles and vesicles are elongated by addition of membranes from various cellular sources, including endosomes, the Golgi apparatus, and other membranous structures (Noda, Suzuki, and Ohsumi 2002; Young et al. 2006). ATG9B is a vertebrate-specific homolog of yeast Atg9, expressed in the placenta, while ATG9A is expressed ubiquitously (Klionsky et al. n.d.; Yamada et al. 2005). ATG9B can compensate for ATG9A in autophagosome formation when ATG9A is suppressed upon autophagy induction (Chiduza et al. 2023; Yamada et al. 2005).

ATG9A and ATG9B function as lipid scramblases, facilitating the bidirectional movement of phospholipids across membranes (Guardia et al. 2020, Chiduza et al. 2023). Recent structural studies have revealed that yeast ATG9, mammalian ATG9A and ATG9B form homo-trimeric complexes, which are essential for their lipid scramblase activity and interaction with ATG2A, a protein involved in membrane delivery (Chiduza et al. 2023; Guardia et al. 2020; Matoba et al. 2020). This trimeric structure is critical for the elongation of phagophores during autophagosome formation. While ATG9A and ATG9B share similar functions, ATG9B is characterized by a longer amino acid sequence, an N-terminal extension, and extended C-termini compared to ATG9A. The C-termini of the homo-trimer forms a HINGE region similar to ATG9A, forming a platform. This platform blocks the cytosolic openings of the central pore and its perpendicular branches. The amino acids in the C-terminal domain of ATG9B are evolutionarily conserved according to the ConSurf database suggesting a critical functional role (Ashkenazy et al. 2016) (supplementary figure 1). Additionally, in mammals, ATG9B contains a longer N-terminal domain with compositional bias with proline-rich and polar-rich residues suggesting a yet-to-be-determined functional role. Moreover, like ATG9A, ATG9B has the tyrosine-based sorting motif suggesting interaction with adapter proteins in its trafficking, which needs further elucidation (Guardia et al. 2020, Chiduza et al. 2023).

Autophagy is crucial in the placental development and trophoblast function under hypoxia (Saito and Nakashima 2013). The placenta is a transient feto-maternal organ facilitating the exchange of gases, nutrients, and waste during fetal development. Cytotrophoblasts are the primary cell type forming the placenta. Upon contact with the maternal endometrium, cytotrophoblast cells fuse to form multinucleated syncytiotrophoblasts (Turco and Moffett 2019). Syncytiotrophoblasts infiltrate the extracellular matrix of the endometrium to enable implantation. Cytotrophoblasts express ATG9B moderately and upon syncytiotrophoblast differentiation, ATG9B expression increases dramatically (Thul et al. 2017) (supplementary figure 2). Under normal conditions, hypoxia stimulates autophagy during placental development. Autophagy is crucial for several trophoblast functions, including degradation of extracellular matrix, infiltration to the endometrium, and migration. Abnormalities in the autophagy process disrupt fetal and placental development resulting in gynecological disorders such as preeclampsia (Hung et al. 2013; Saito and Nakashima 2013). Autophagy is also essential for embryonic development; various genetically modified mouse models have been generated to study the functions of autophagy genes in mice. The knockout of the core autophagy genes has resulted in embryonic death (*Becn1, Pik3c3/Vps34, Atg9a, Rb1cc1, Atg13*) or neonatal death (*Ulk1/2, Atg3, Atg5, Atg7, Atg12, Atg16l1*) (Kuma, Komatsu, and Mizushima 2017). Conditional knockout of *Atg7* in trophoblasts led to the inhibition of autophagy, reduced trophoblast invasion, increased apoptosis, anomalies in vascular remodeling, and overall inefficient placentation, underscoring the importance of autophagy in placental development (Aoki et al. 2018). In studies with Atg9 genes, conditional knockout of *Atg9a* in mouse brain tissue resulted in neonatal death in 50% of the mice. None of the animals lived longer than four weeks after birth. In somatic neurons SQSTM1/p62 and NBR1 receptor proteins and ubiquitin were accumulated, suggesting impaired autophagy at day 15. At day 28, the level of these proteins was reduced suggesting that autophagy was functional to some extent. Atg9a knockout also caused the axonal degeneration of Purkinje cells in the cerebellum, defects in neuronal circuit formation, and dysgenesis of the corpus callosum. Additionally, cultured primary cells showed impairment of the neurite extension in neurons (Yamaguchi et al. 2018).

These studies stress the importance of autophagy in placental and fetal development and neurogenesis in humans and mice. Impairment of autophagy and knockout of core autophagy genes can result in gestational problems, fetal growth restriction, or death.

Autophagy is a core cellular degradation mechanism, its perturbations are often linked with human diseases, including but not limited to mutations in core autophagy genes. Mutations and expressional variations of autophagy genes are often linked with neurodevelopmental and neurodegenerative diseases and various cancers (Yang and Klionsky 2020). Mendelian diseases associated with *ATG5* and *ATG7* mutations have been linked to spinocerebellar ataxia. (M. Kim et al. 2016). *ATG7* mutations were also associated with neurodevelopmental disorders. The patients showed similar symptoms of ataxia, developmental delay, optic atrophy, bilateral sensorineural hearing loss, spastic paraplegia, and facial dysmorphism (Collier et al. 2021). Both gene mutations led to impaired autophagy flux.

*ATG9B* variants are not reported in the literature as pathogenic for Mendelian diseases however, frameshift mutations in ATG9B were reported in 13 different types of cancer, including breast, hepatocellular, gastric, and colorectal cancers and coronary artery disease. These mutations disrupt normal autophagic processes, underscoring the gene’s pivotal role in many diseases (Kang et al. 2009; Mehrabi Pour et al. 2019). The *ATG9A* gene is not associated with an inherited genetic disease, but similar to *ATG9B* it was linked to triple-negative breast cancer (TNBC) and invasive ductal carcinoma (IDC) cancer pathology. In TNBC, ATG9A is overexpressed in transcript and protein levels in cancer tissues compared to healthy adjacent tissues. (Claude-Taupin et al. 2018). ATG9A downregulation was linked to trastuzumab-resistant cells, leading to Her2 escape from lysosomal degradation (Nunes et al. 2016).

Our study has originated from the Blue Gene project, a screening project for pediatric patients with neurodevelopmental disorders, born to consanguineously married couples. In the Blue Gene project, whole exome sequencing (WES) was adopted to identify novel candidate gene variants. In this manuscript, we describe a novel, rare genetic neurodevelopmental disorder caused by an eleven-nucleotide deletion in the *ATG9B* gene. This frameshift mutation in ATG9B gene exon 9 causes the alteration of six amino acids (695-700) and introduces a premature stop codon at the 701^st^ position truncating the ATG9B protein. The two affected subjects were born to a consanguineously married couple and exhibited mental retardation, facial dysmorphia, obesity, and attention deficit. The index patient’s younger sister presented with similar but less severe symptoms. The siblings were homozygous for the mutation while the parents were heterozygous. We characterized the mutation by *in vitro* (expression in mammalian cells) and *in vivo* (genetically engineered mouse models) methods to study this rare genetic disorder.

## Materials and Methods

### Identification and validation of candidate gene variant

The index case in this work was identified through the Blue Gene project where patients and their families were screened for novel candidate gene variants (Ethics committee approval Protocol no: 446-SBKAEK) a sample of informed consent forms is presented in supplementary file 1). Peripheral blood samples from the family were collected, and genomic DNA was isolated using the Thermo Scientific Pure-Link Genomic DNA Mini Kit (K182001). The WES was analyzed as previously described (Hiz et al. 2022). The analysis revealed a homozygous 11-nt deletion in the *ATG9B* gene of affected siblings while parents were heterozygous for the deletion, suggesting a Mendelian disease. The deletion was validated by Sanger sequencing of the PCR amplified mutation site in the index case and by agarose gel electrophoretic analysis in other family members. Primers used for amplification of patient genomic DNA for the variant verification by Sanger sequencing are listed in the supplementary table 1 (Primer 1, 2).

### Cloning of the WT and truncated ATG9B

The human ATG9B coding sequence (NCBI Reference Sequence: NM_001317056.2) was synthesized commercially, sequence was verified by the depositor and after cloning by us. An N-terminal FLAG tag was added by PCR and the resultant construct was cloned into pcDNA3.1. To generate the mutant ATG9B, we added six amino acids altered after the frameshift and added a stop codon at position 701. Amplification of the constructs, addition of FLAG tag and altered amino acids for truncated were achieved by two sequential PCRs (Primers 3-7). Constructs were subsequently subcloned into 3xFLAG CMV 10 plasmid. To generate ATG9B WT-myc and ATG9B TR-myc constructs the sequence was subcloned into pcDNA3.1 myc his A. These plasmids were free of FLAG tag (Primer 8). For the generation of HeLa stable cells, the FLAG ATG9B WT construct was subcloned to pBOBI lentiviral plasmid (Primer 9). HeLa cells were infected and monoclonal stable HeLa cells expressing ATG9B WT (HeLa ATG9B WT clone) were obtained (Primers used for cloning are listed in the supplementary table 1).

### Transfection, Western blotting, RT-PCR

HEK293T and HeLa cells were transfected using Promega Fugene HD. Protein extracts from transfected cells were analyzed by western blotting using commercial primary antibodies (Sigma anti-FLAG M2 F1804, Santa Cruz anti-GFP SC9996, Abcam Anti-β Actin ab6276).

For RT-PCR assays, total RNA extraction from cells or mouse tissues was performed using the MN NucleoSpin RNA Mini kit (740955.50) according to the manufacturer’s protocol. cDNA synthesis was performed with Thermo Scientific RevertAid cDNA First Strand Synthesis kit (K1621). RT-PCR primers used in mouse placenta samples are listed in supplementary table 1.

### Immunocytochemistry and immunohistochemistry

HEK293T and HeLa cells were transfected with ATG9B WT and truncated (TR) constructs. 24 hours post-transfection, immunofluorescence was performed. Primary antibody (Sigma anti-FLAG M2 F1804) incubation was overnight at 4°C and secondary antibody (Cell Signaling Alexa Fluor 594 anti-mouse 8890S, and 488 anti-rabbit 4411S) was 1 hour at RT. For colocalization imaging in Figure 3, Zeiss LSM 800 confocal microscopy was used. Fluorescence pictures were captured with Olympus Upright BX61 microscope and colocalization was analyzed by FiJi (Figure 4) (Schindelin et al. 2012).

For immunohistochemistry studies, human placenta tissues were fixed in 4% formaldehyde immediately after isolation. Subsequently, sections were stained by IHC, using a home-made anti-ATG9B rabbit polyclonal primary antibody and ScyTek Laboratories SensiTek HRP (Anti-polyvalent) kit.

Histomorphometry analysis of the mouse placenta was performed on 18.5 dpc embryos isolated from WT/KI female and KI/KI male crosses. Litter in each conceptus were genotyped by the total DNA isolated from embryonic tail tissue. Placentas were fixed, processed, and embedded in the paraffin. Hematoxylin-eosin staining was performed on 4 µM sections. The decidua, junctional, and labyrinth zone widths were measured at six different points for subsequent histomorphometry analysis.

### Generation of Atg9b knock-in mice

Single guide RNAs (sgRNAs) targeting the mutation site in the mouse Atg9b locus were selected based on their proximity to the mutation site, on-target efficacy, and minimal off-target activity. Single-stranded oligodeoxynucleotides (ssODNs), synthesized to extremer quality by Eurofins, were used as templates for homology-directed repair (HDR) (supplementary table 1 -sgRNA 1, 2, HDR template). sgRNAs were synthesized using the NEB HiScribe T7 High Yield RNA Synthesis Kit (E2040L) and then purified using the Monarch RNA cleanup kit (T2040L). The Cas9/sgRNA complexes and ssODN oligos were introduced to E0.5 stage mouse embryos by electroporation. The embryos were transferred to pseudopregnant CD1 females (Modzelewski et al. 2018).

For genotyping of newborn pups, tail biopsies were prepared according to the protocols described previously (Erguven et al. 2023). PCR and restriction digestion were employed to verify HDR-mediated alterations (supplementary table 1, Primer 10, 11). The genotyping strategy involved replacing the StuI restriction site in the wild-type sequence with an EcoRI site by HDR template. Successful alterations were confirmed by restriction digestion and Sanger sequencing.

### Behavioral tests

Given that the most prominent symptom in the patients was intellectual disability, a comprehensive series of memory tests were designed (Bailey, Kandel, and Harris 2015; Kandel, Dudai, and Mayford 2014). Perirhinal cortex-dependent memory was evaluated using the novel object recognition (NOR) test, while hippocampus-dependent memory was assessed through the novel location recognition test (NLR). Social memory, which primarily involves the hippocampus, amygdala, and prefrontal cortex, was measured using the social novelty test. Amygdala-dependent memory was evaluated by the passive avoidance test. In addition to memory assessments, other behaviors commonly affected in neurodevelopmental disorders were evaluated. Stereotypic movements, social interaction, and anxiety-like behaviors were evaluated by marble burying test, social preference test, and the open field test, respectively (Brown, Stanford, and Schellinck 2000; Eltokhi, Kurpiers, and Pitzer 2020). The tests were performed according to standardized protocols in the literature.

SPSS (Statistical Package for the Social Sciences) was used for the statistical analysis of all data. Normally distributed data, namely total distance (cm) and total time spent in the center area (sec) in OFT, buried marbles, social preference, NOR, NLR and social novelty scores were analyzed by Student t-test. Latency to first enter the center area (sec) in OFT was analyzed by Mann-Whitney U-test. Passive Avoidance data was analyzed by Kaplan-Meier Survival Analysis. Data are presented by the mean ± Standard Error of Mean (SEM).

## Results

### Case report

The Blue Gene project aimed to identify candidate gene variants causing Mendelian diseases in pediatric patients with neurodevelopmental disorders. Genomic DNA samples from children of consanguineous marriages and their parents were analyzed by WES, with consent. The index case, a 12-year-old male patient (II-1) presented with intellectual disability. His medical history revealed delayed walking, inability to read and write, and limited speech. Due to his intellectual impairment, he was attending a special education program. There was a history of first-cousin marriage between his parents. The family had two other children besides the index case: a 9-year-old sister (II-2) with milder intellectual disability and obesity who was also receiving special education, like the index case. Their 16.5-year-old brother had severe motor and intellectual disability, was unable to walk or talk, experienced epileptic seizures and hearing loss.

On physical examination, the patient weighed 72 kg (>2SD), had a height of 153 cm (50^th^ percentile), and a head circumference of 54 cm (25^th^ percentile). He made eye contact and had a good-natured, affectionate temperament. He followed commands but had very limited verbal communication, giving short, one-word answers. He was frequently distracted and had difficulty concentrating during the examination. He was obese, had deep-seated eyes, and had a Simian line on his right hand. Other systemic and neurological examinations were normal. Biochemical, metabolic, and hormonal tests showed no abnormal findings. His cranial MRI was normal.

### Identification of the ATG9B mutation

The patient (II-1), his sister (II-2), and his parents (I-1, I-2) genome were subjected to WES. Their older sibling could not participate in genetic testing (Figure 1A). WES readings revealed an 11-nucleotide deletion in exon 9 of the ATG9B gene (Figure 1B). The variant rs747858674 is positioned at NM_173681.5 (ATG9B):c.2083_2093del (p.Leu695fs). The mutation, not linked to any clinical pathogeny, had a global frequency of 0.00008640 (139 out of 1608812 alleles) according to gnomAD (Chen et al. 2023). The highest frequency is observed in Middle Eastern populations at 0.0008278 (5 out of 6040 alleles) followed by Ashkenazi Jewish population 0.0005781 (17 out of 29406 alleles). Sanger sequencing confirmed the deletion for the proband. Separation of the alleles on the agarose gel electrophoresis confirmed the mutation for the family members. The affected siblings were homozygous for the deletion, while the parents were heterozygous (Figure 1C).

**Figure 1.**
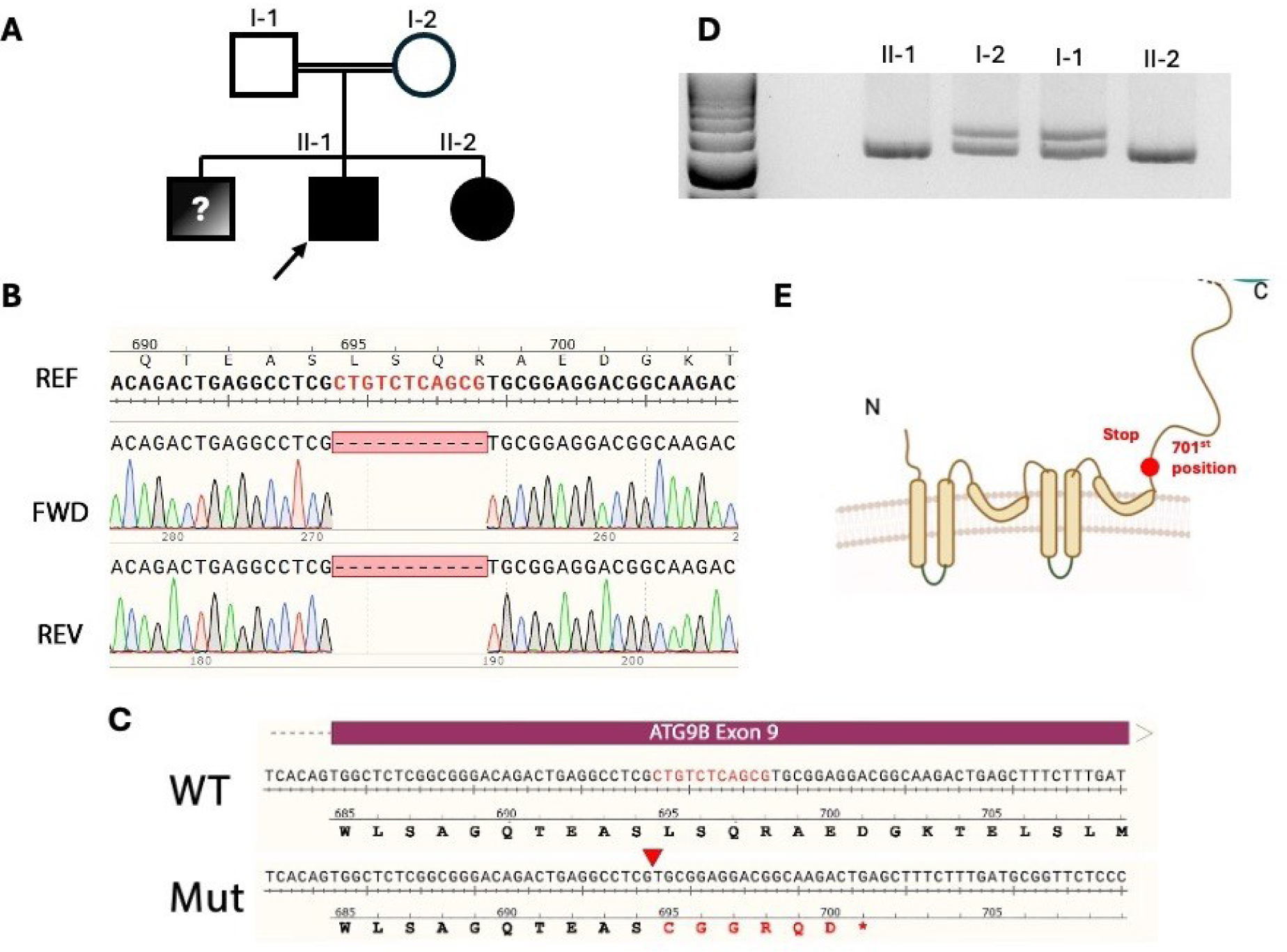
Identification and validation of the *ATG9B* mutation. A) Family tree of the pediatric patient is depicted. Parents (I-1 and I-2) are first cousins and asymptomatic. “?” indicates the firstborn male of the family with motor-mental disorder and epilepsy in addition to intellectual disability. He was not involved in this study. The proband (II-1) is the second-born male presents with intellectual disability, limited speech, attention deficit, facial dysmorphia, and obesity. The youngest female child (II-2) showed similar symptoms as the proband but milder. B) 11-nt deletion was detected by WES at the exon 9 of the *ATG9B* gene for the proband and younger sister. The parents were heterozygous. Sanger sequencing confirmed the deletion for the proband C) The deletion causes a frameshift and change of 6 amino acids then changes aspartic acid at position 701 to a stop codon, truncating ATG9B protein. D) The deletion alleles in the family were confirmed by agarose gel electrophoresis. 11-nt deletion causes the formation of shorter DNA fragment which can be distinguished from the WT allele after electrophoretic separation. The affected children are homozygous for the deletion.

The 11-nt deletion causes a frameshift and alteration of six amino acids, followed by addition of a premature stop codon (p.Leu695fs). This frameshift deletes the C-terminal cytosolic domain of the ATG9B protein, resulting in a shorter, truncated ATG9B (Figure 1D, 1E). This variant is not associated with any known genetic disease, and no clinical significance is reported in the ClinVar database (Landrum et al. 2018). In this work, we propose this deletion is a disease-causing mutation with autosomal recessive inheritance.

### ATG9B expression increases upon syncytialization

Although ATG9A is expressed ubiquitously, ATG9B expression is limited to the placenta. It is moderately expressed in cytotrophoblast cells and highly expressed in syncytiotrophoblast cells (Supplementary figure 1A,B) (Thul et al. 2017; Yamada et al. 2005). Syncytiotrophoblasts are essential for embryonic development and functional placenta (Soares, Iqbal, and Kozai 2017). Placenta-derived BeWo and JAR cells are choriocarcinoma cells with trophoblast origin. Moreover, BeWo cells constitute a good placenta syncytiotrophoblast model since they exhibit cell fusion upon forskolin treatment and express human chorionic gonadotropin β (hCGβ), a syncytiotrophoblast marker (Orendi et al. 2010). By contrast, JAR cells respond to treatment and express hCGβ, but they do not fuse failing to serve as a syncytialization model (Li et al. 2023). Therefore, we induced BeWo and JAR cells with forskolin for 48 hours to allow syncytialization. After 48 hours, the cell membranes of BeWo fused, as expected (Figure 2A). hCGβ expression increased in both cell lines, however ATG9B expression increased only in BeWo cells (Figure 2B).

**Figure 2.**
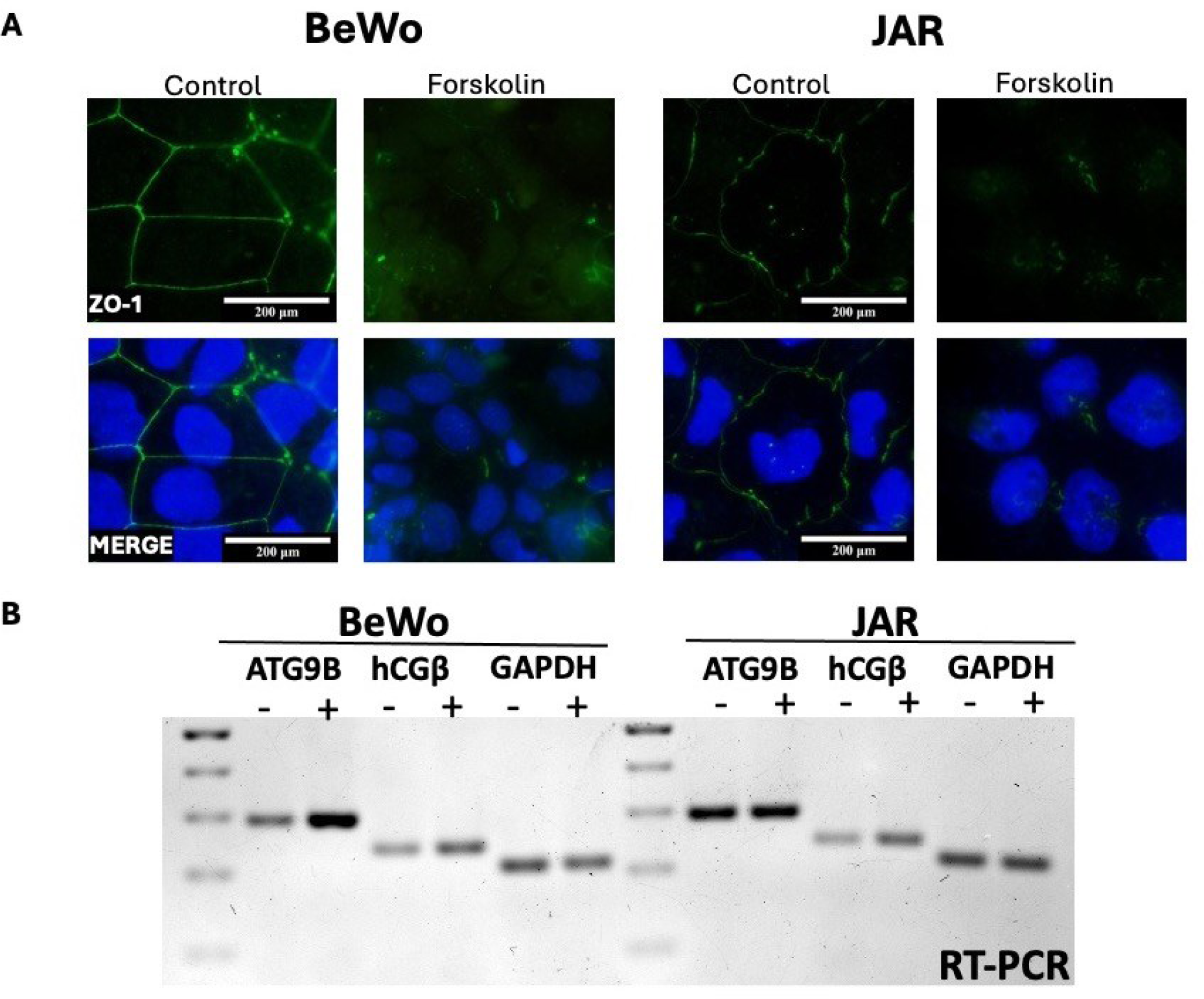
Increase in ATG9B expression upon forskolin-induced syncytialization. A) Placenta-derived BeWo and JAR cells were treated with forskolin for 48 hours, and immunofluorescence staining was performed with tight junction protein Zona Occludens-1 (ZO-1) to determine the cell boundaries. The control condition is treated with DMSO. The cell membranes fused upon forskolin treatment, suggesting syncytialization. B) Upon forskolin treatment (+), hCGβ syncytiotrophoblast marker expression increased in both cell lines at the RNA level. In BeWo cells, ATG9B expression increased in parallel, while GAPDH levels were similar.

### Localization of WT and truncated ATG9B in cells

ATG9A localizes in Golgi and trans-Golgi vesicles, and ATG9B was recently shown to localize to Golgi (Chiduza et al. 2023; Young et al. 2006). Therefore, we aimed to determine the localization of WT and truncated ATG9B by immunofluorescence microscopy analysis. We transiently transfected HeLa cells with ATG9B WT and TR constructs. They both localized to trans-Golgi vesicles marked by Golgin-97. ATG9B TR localization was aberrant large puncta in perinuclear sites and caused abnormal morphology in Golgi vesicles similar to Golgi fragmentation. Additionally, we transfected a stable HeLa monoclonal cell line expressing FLAG-tagged full length human ATG9B, with myc-tagged WT and truncated ATG9B and analyzed their colocalization. We observed that ATG9B TR colocalized with WT protein, albeit on abnormal vesicular structures which appear only upon ATG9B TR expression.

### Truncated ATG9B is not stable

We have carried out experiments to determine and compare the subcellular localization of WT and mutant ATG9B proteins. When we transfected mammalian cells for ectopic expression and subsequent immunofluorescence microscopy analysis, we frequently observed that ATG9B TR expression is scarce in comparison to WT. This could be because of RNA or protein instability. To address this, we designed a controlled experiment where equal amounts of WT and truncated ATG9B constructs were transfected in human cell lines. Analysis of western blot results from protein extracts showed that ATG9B TR expression was significantly lower than WT. This was confirmed by immunofluorescence (Figure 3A, 3B). These results suggest that truncated ATG9B protein is not stable when expressed in cells, as RNA levels were similar between WT and TR (Figure 3C).

**Figure 3.**
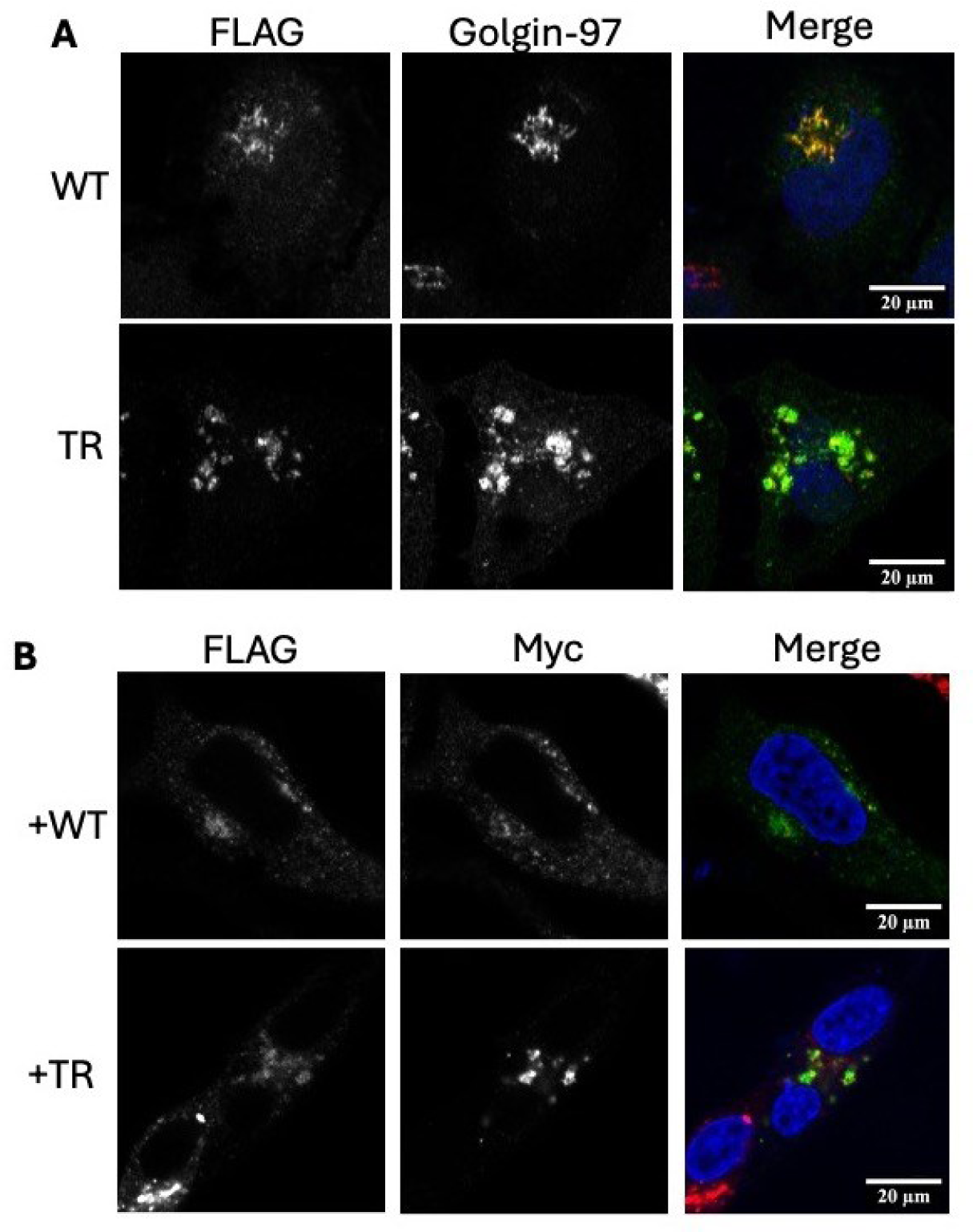
Colocalization analysis with ATG9B WT and truncated forms. A) HeLa cells were transiently transfected with FLAG ATG9B WT or TR constructs. Immunofluorescence was performed with anti-FLAG and anti-Golgin-97 antibodies. Golgin-97 is a trans-Golgi marker and endogenous protein was detected. ATG9B WT and ATG9B TR both localize to trans-Golgi compartments, while truncated protein causes abnormal Golgi structure. B) HeLa ATG9B WT clone was transfected with ATG9B WT-myc or ATG9B TR-myc constructs. ATG9B TR colocalizes with the WT on abnormal membranous structures.

### Development of a knock-in mouse model expressing truncated Atg9b

Genetically engineered mouse models are well-established tools for modeling human diseases. Therefore, we generated a mouse model for the human disease resulting from the ATG9B mutation. We first analyzed the human and mouse protein sequences. C-terminal cytosolic domains are evolutionarily conserved (88% identity) suggesting a critical function (Figure 4A). We designed two sgRNAs targeting the mutation region in exon 9 of the mouse Atg9b locus. We used an HDR template to introduce a STOP codon at 694^th^ position of the mouse *Atg9b* gene. Simultaneously, we disrupted the 5’ PAM sequence and changed the StuI restriction site to EcoRI. The C-terminal sequence is thereby truncated. C57B6/J WT mouse embryos were electroporated with the Cas9/sgRNA complex and ssODNs, and subsequently transferred to pseudo-pregnant CD1 females to generate founder mice (Figure 4B). For genotyping, we amplified the locus by PCR and performed restriction digestion to distinguish WT and knock-in (KI) alleles (Figure 4C). Lastly, we confirmed the knockin allele by sequencing (Figure 4D).

**Figure 4.**
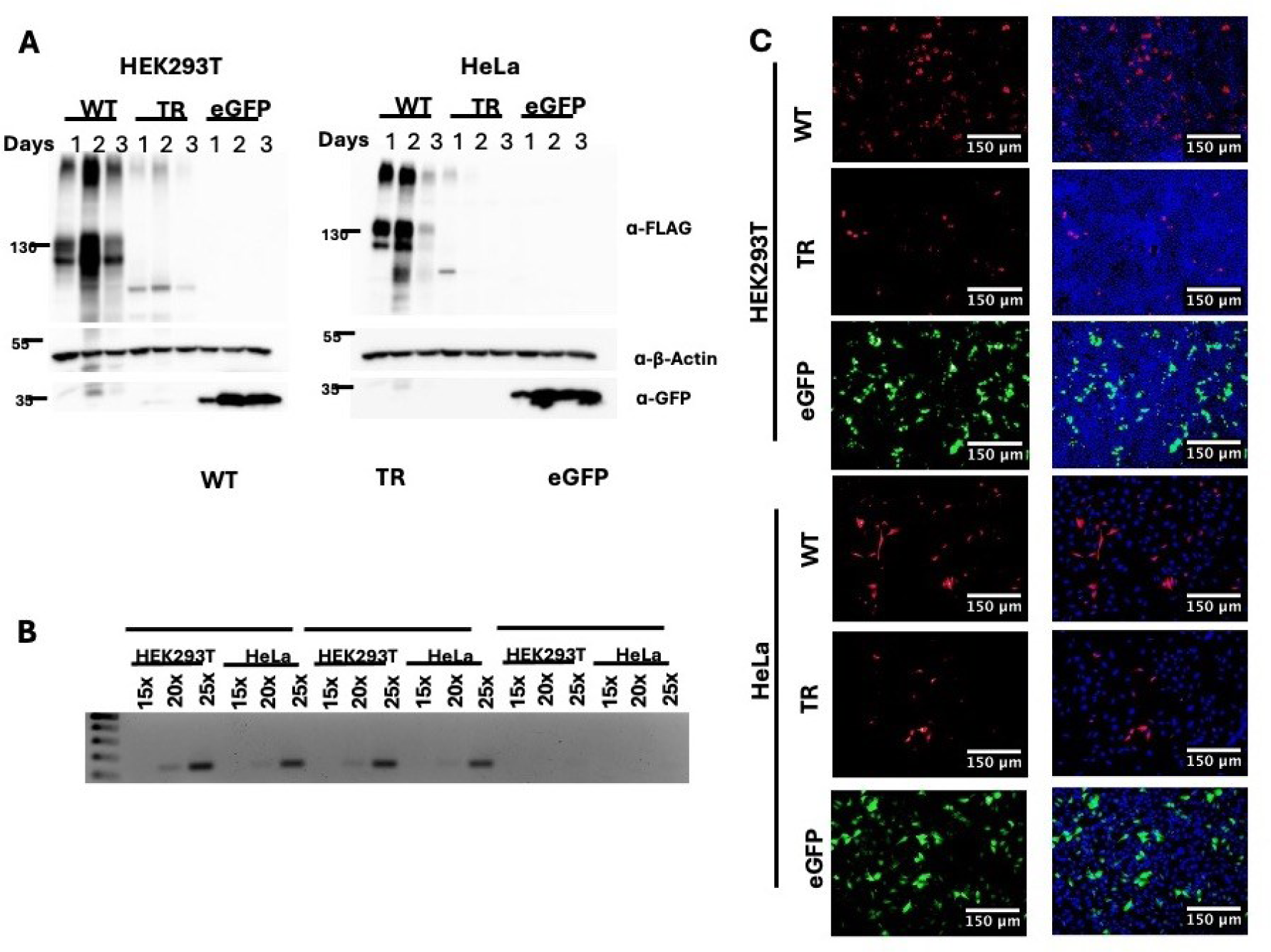
Truncated ATG9B protein is not stable. A) HeLa and HEK293T cell lines were transfected with WT and truncated ATG9B constructs. eGFP was used as a control. Cell pellets were collected for western blot and coverslips were fixed for immunofluorescence staining on post-transfection day 1, day 2, and day 3. WT ATG9B expression is detected on all three days, the highest at day 2 for both cell lines. Truncated ATG9B expression was minimal in comparison. B) On day 3, cell pellets were obtained for assessing transcriptional expression level. The WT and truncated ATG9B RNA levels were comparable. C) Representative images of immunofluorescence experiment confirming the western blot analysis.

Homozygous knockin mice were viable, had no obvious growth abnormality at birth and in the neonatal period, they reached adulthood and were fertile. Mating heterozygous males and females yielded offspring numbers expected from a Mendelian distribution pattern of a non-pathogenic allele (WT/WT:30, WT/KI:42, KI/KI:28).

### Histological observations in human and mouse placenta

Human ATG9B is specifically expressed in the placenta, especially in syncytiotrophoblasts (Yamada et al. 2005; Thul et al. 2017; Uhlén et al. 2015). In our cell culture syncytiotrophoblast model, we detected increased ATG9B upon induced syncytialization at the RNA level. To address ATG9B protein expression in human term placenta, tissue samples were obtained immediately after delivery for immunohistochemistry analysis. Using a homemade rabbit polyclonal antibody targeting ATG9B N-terminal domain (amino acids 134-207), we detected a moderate expression of ATG9B in cytotrophoblasts (Figure 5A, blue arrowhead) and more prominently in syncytiotrophoblasts (Figure 5A, black arrowhead). This study provides the first demonstration of ATG9B expression in the placenta at protein level.

**Figure 5.**
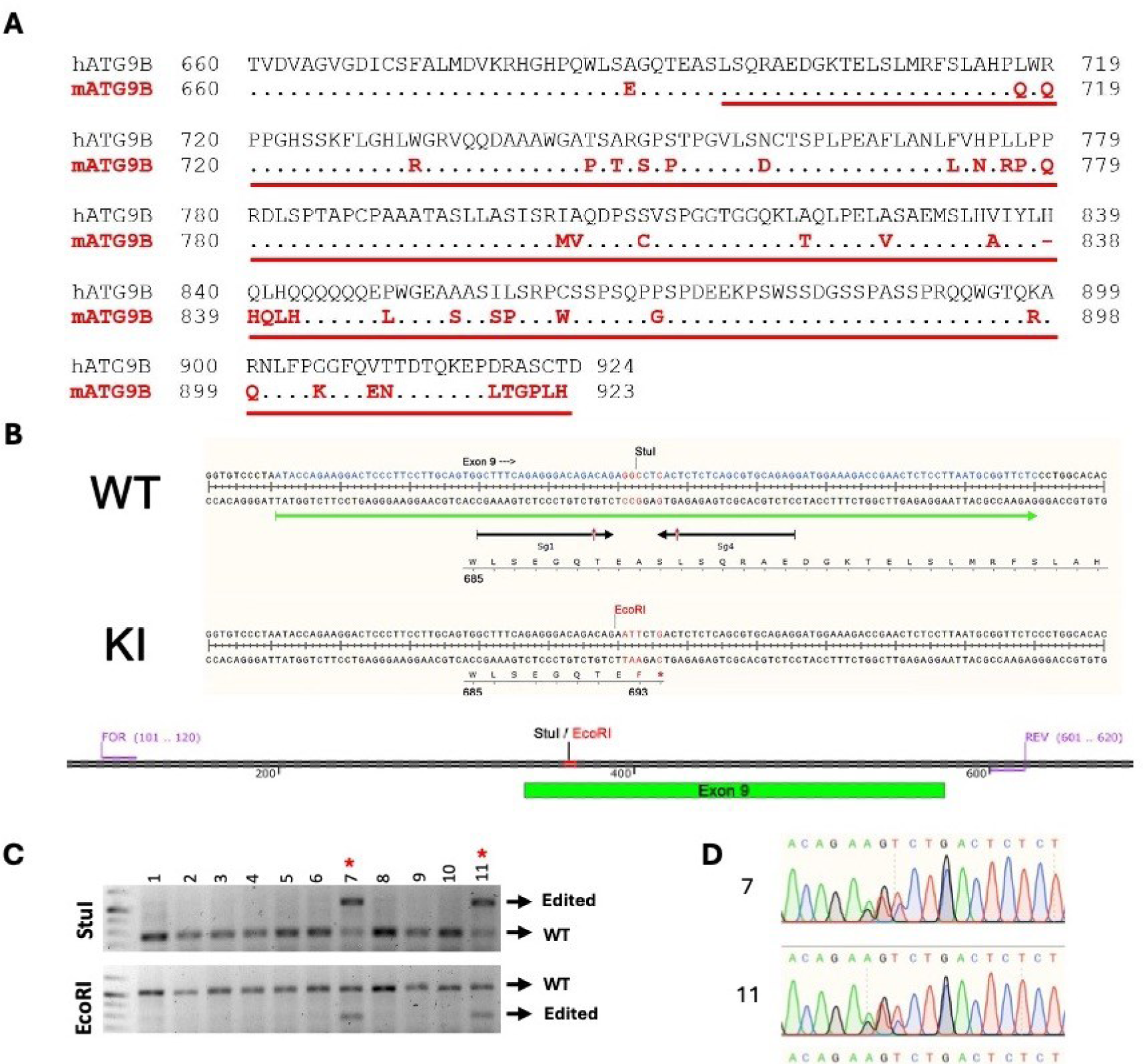
The generation of Atg9b knock-in mouse model. A) The alignment of the C-terminal domains of human and mouse ATG9B indicating strong conservation. B) We designed two sgRNAs to target the mutation site in Exon 9 for Cas9-mediated double-strand break. HDR template was designed to introduce a stop codon, change StuI site to EcoRI, and destroy PAM sequence of the sgRNA1. Alanine to phenylalanine alteration was a consequence of the desired alterations. C) The genotyping strategy of knock-in mice involved PCR and restriction digestion with StuI and EcoRI to determine both WT and knock-in alleles. Heterozygous mice 7 and 11 were selected for Sanger sequencing confirmation. D) Sequence validation of the heterozygous mice confrimed the desired alteration. The double peaks in the chromatogram indicate the occurrence of both WT and edited alleles in heterozygous mice. These mice were selected for breeding.

**Figure 6.**
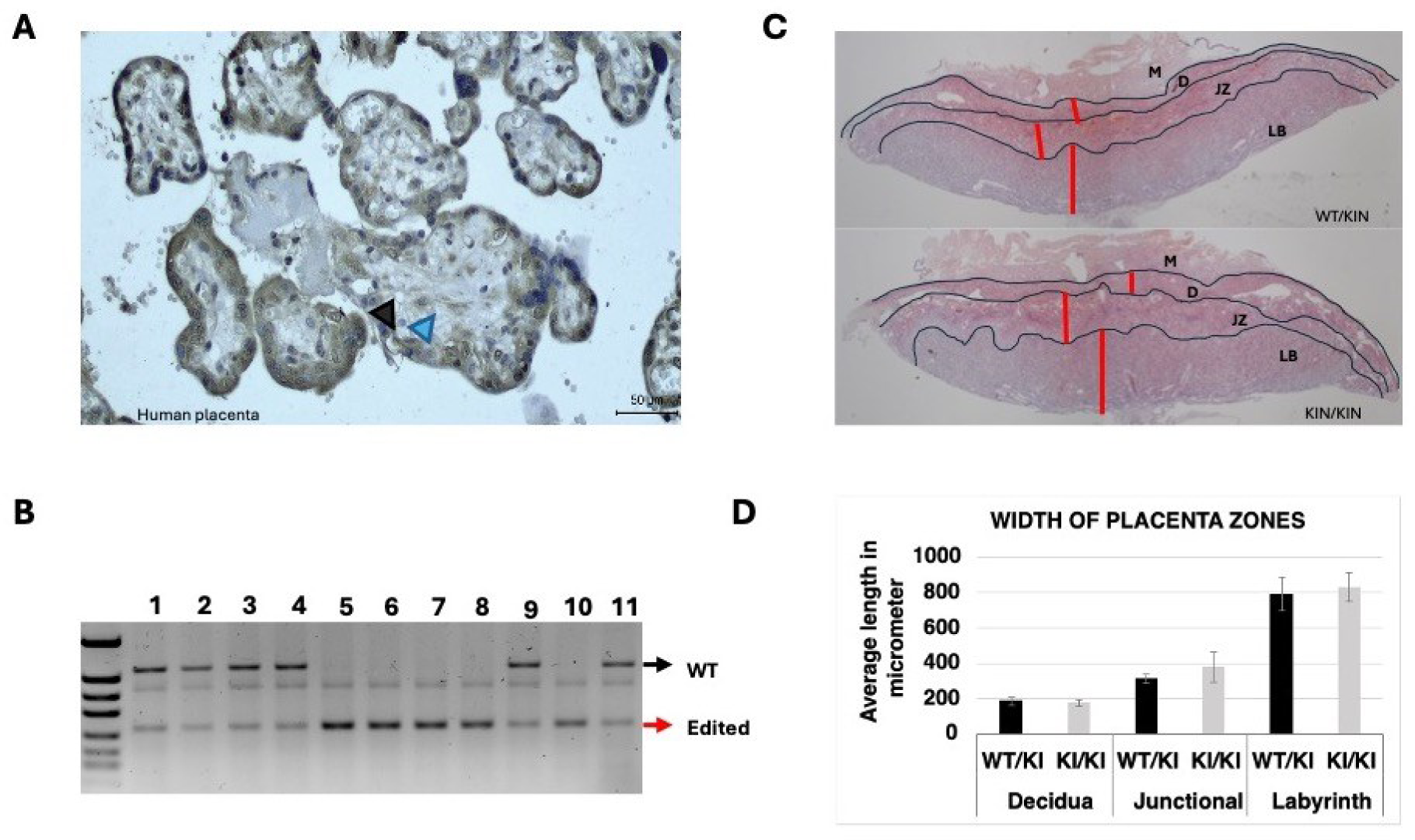
Human and mouse placental examinations. A) Immunohistochemistry staining by home-made ATG9B antibody shows high expression of ATG9B protein in syncytiotrophoblast (black arrow-head) and moderate expression in cytotrophoblasts (blue arrow-head). B) Timed-breeding was performed with WT/knock-in females and knock-in/knock-in males. For placental examinations with litter-mate mouse conceptuses, hotshot lysates from the embryonic tissues obtained at 18.5 dpc were amplified and digested with EcoRI. Conceptus 1, 2, 3, 4, and 9 were heterozygous, and 5, 6, 7, 8, and 10 were homozygous knock-in. C) Representative image of placenta at 18.5 dpc showing matrial gland (M), Decidua (D), Junctional zone (JZ) and Labyrinth (L). The zones were marked by black line. Red lines indicate a single width measurement, for each zone 6 measurements were obtained, and the average was calculated. D) Graphical representation of the average measurements in micrometers. The heterozygous (WT/KI) and homozygous (KI/KI) widths were analyzed by unpaired student’s T-test and no statistically significant difference was detected in decidua, junctional zone or labyrinth.

**Figure 7.**
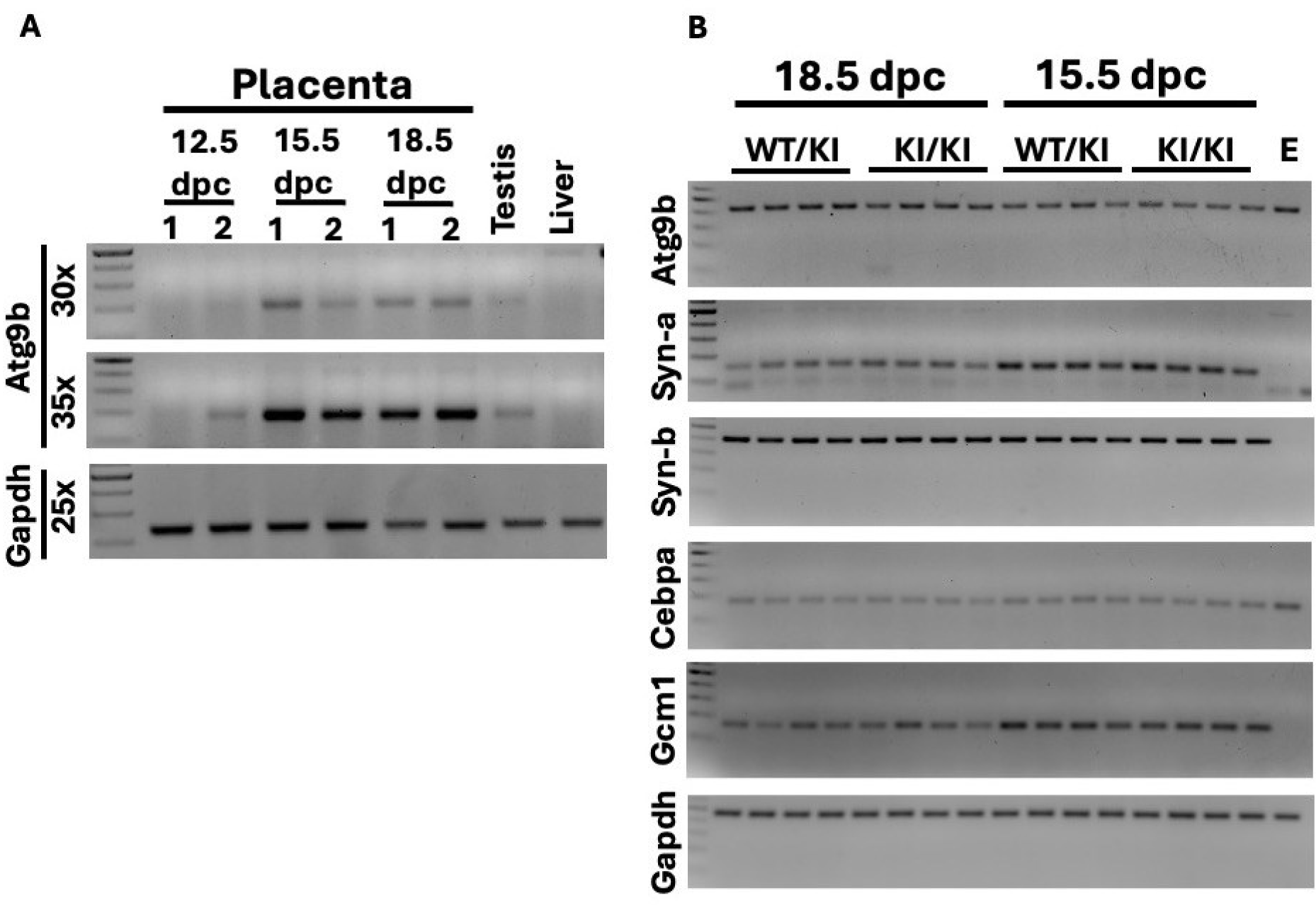
Transcriptional analysis of Atg9b and genes important for placental development. A) Timed-breeding was performed with WT mice and placentas were isolated at 12.5 dpc, 15.5 dpc 18.5 dpc developmental points. RT-PCR was set with a duplicate mix for 30 and 35 cycles. As control Gapdh was amplified for 25 cycles. The Atg9b expression at 12.5 dpc was low, and only detected after 35 cycles compared to 15.5 and 18.5 dpc placenta detected by 30 cycles. Atg9b was detected moderately in testis but not in liver. B) Timed-breeding was performed with WT/knock-in females and knock-in/knock-in males. 4 WT/KI and 4 KI/KI per stage were subjected to RT-PCR. Atg9b expression was comparable in both WT/KI and KI/KI placenta samples in both stages. Syn-a expression was higher at 15.5 dpc and Synt-II marker Syn-b expression was similar in both developmental points. Cebpa and Gcm1 expression was higher at 15.5 dpc. The expression level of the genes involved in trophoblast and syncytiotrophoblast development was not affected by Atg9b knock-in in either stage. The Gapdh control levels were similar.

Neurodevelopment in the fetus could be affected from placental abnormalities during pregnancy Therefore, we investigated the placental structure in homozygous knock-in and heterozygous control mouse placentas at 18.5 dpc developmental stage (Rosenfeld 2021). Tissues from littermate embryos were compared by histomorphometry analysis. Placental samples were isolated from homozygous knock-in male and heterozygous female intercrosses and subsequently genotyped (Figure 5B). Hematoxylin-eosin staining was performed on midsections of the placenta marked by the umbilical cord, showing three major zones: decidua, junctional, and labyrinth (Figure 5C). Six width measurements for each zone were randomly taken from each placenta. An unpaired T-test between the two groups indicated no significant differences in the structure between heterozygous and homozygous placentas (Figure 5D). Thus, histomorphometry analysis did not reveal abnormalities in placentas from homozygous mutant embryos. Furthermore, no developmental delay in homozygous mouse embryos was observed (data not shown).

### Comparison of gene expression

We used RT-PCR to examine expression differences in *Atg9b*, and other genes involved in syncytiotrophoblast formation. Syncytiotrophoblasts differentiate and circle maternal blood vessels by 9.5 dpc. By 15.5 dpc syncytiotrophoblast I and II layers protrude into the cytotrophoblast layer and start releasing hormones into maternal blood. Day 18.5 dpc is human-term placenta equivalent in C57B6/J mouse (Elmore et al. 2022; Shi et al. 2004). 18.5 and 15.5 dpc heterozygous and homozygous knock-in placentas from littermate embryos were isolated for analysis. In addition to *Atg9b*, we analyzed the expressions of *Gcm1, Cebpa, SynA and SynB*. Glial Cells Missing Homolog 1 (*Gcm1*) regulates cytotrophoblast fusion to form syncytiotrophoblast layer (Lu et al. 2013; Simmons et al. 2008). Gcm1 activates Syncyntin-A and Syncyntin-B (SynA and SynB respectively) envelop proteins are expressed in Syn-I and Syn-II cells and are enrolled in cell fusion for the formation of syncytiotrophoblast layers (Lu et al. 2013; Simmons et al. 2008). CCAAT/Enhancer Binding Protein Alpha (Cebpa) is an important transcriptional factor for trophoblast differentiation (Lu et al. 2013; Simmons et al. 2008). We hypothesized that the truncation of *Atg9b* may result in alterations in the expressional pattern of *Gcm1*, *SynA, SynB*, and *Cebpa* due to the abnormalities in development of syncytiotrophoblasts. *Atg9b* expression did not differ significantly between heterozygous and homozygous knock-in placentas., though it was higher at 18.5 dpc compared to 15.5 dpc. However, *SynA and Gcm1* expression was lower at 18.5 dpc, while *SynB* was higher. While the expression of these genes varies between 15.5 and 18.5 dpc placenta samples, their expression levels in homozygous knock-in placentas did not change. Therefore, Atg9b truncation does not affect the expression of genes related to syncytiotrophoblast differentiation in mice.

### Behavioral studies

Pediatric patients harboring homozygous ATG9B mutation presented with intellectual disability which was the most prominent clinical feature. Neurobehavioral tests designed and standardized to assess stereotypic behavior, memory and cognition were employed. These tests are extensively used in the literature in different models including but not limited to genetically modified mice (Van Meer and Raber 2005). Therefore, we applied a series of behavioral tests to WT and homozygous knock-in mice to assess memory and anxiety-like behaviors (Figure 8A).

**Figure 8.**
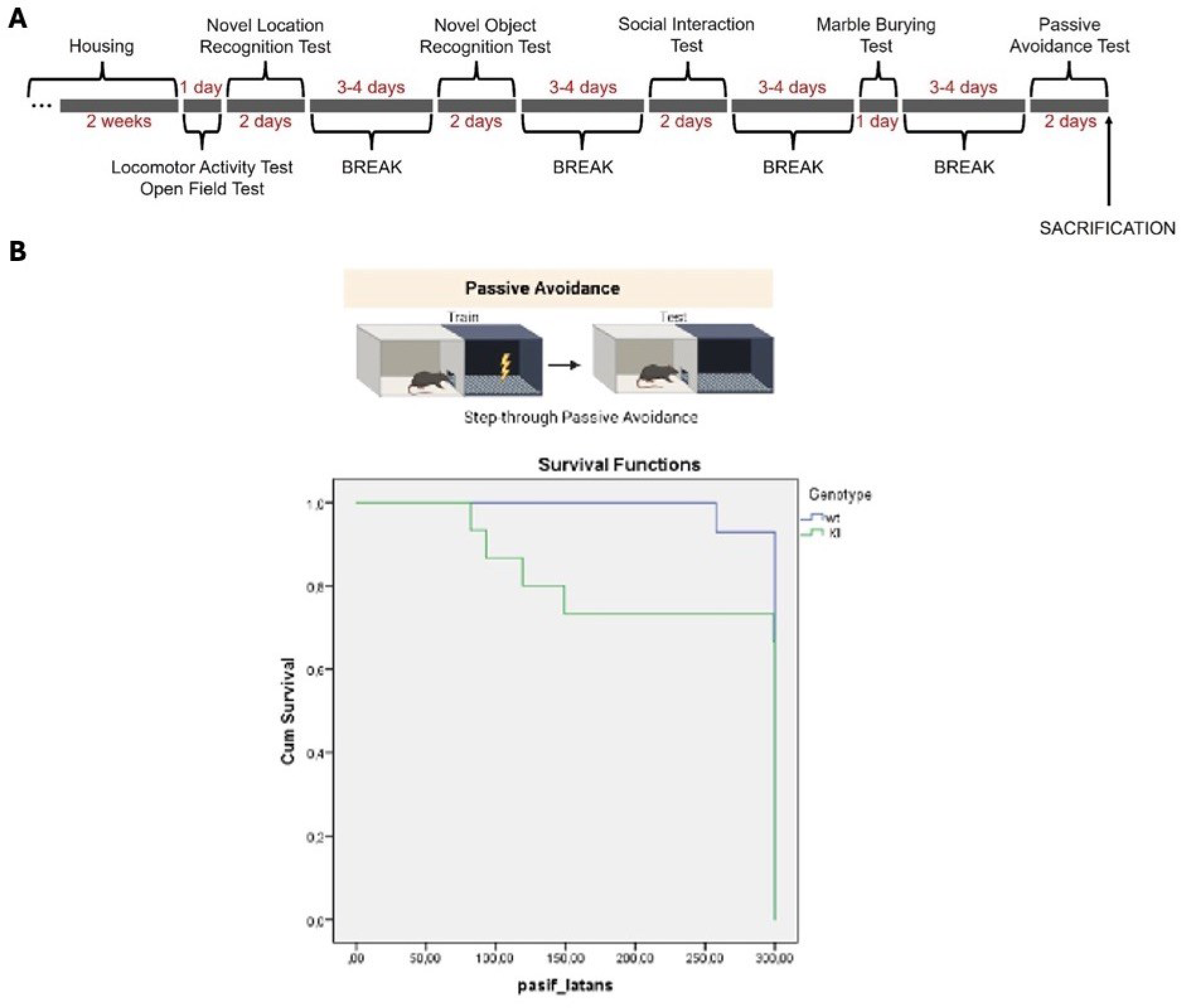
Behavioral assessment of WT/WT and KI/KI adult male mice. A) Experimental plan for the assessment. B) Passive avoidance test was conducted in a cage with one bright compartment and one dark compartment separated by a door. Mouse was placed in the bright compartment and 10 seconds later the door was opened. When the mouse moved to the dark compartment based on their natural inclination, the door closed, and the mouse received a 0.9 mA electric shock for 2 seconds. One hour later, the test was repeated and the latency of mice to enter the dark compartment was recorded. The animals that did not cross into the dark compartment after 5 minutes were returned to their cages and their latencies were recorded as 300 seconds. The analysis of videos were analyzed by Ethovision XT8 program.

Locomotor activity was measured using the open field test, showing no significant differences between WT and ATG9B homozygous KI groups (p=0.950, t=0.63) (Figure S3). Additionally, the time spent in the central area (p=0.208, t=-1.297) and the latency to first enter the central area (p=0.116) were similar between the WT and knock-in groups (Figure S3A), suggesting that the ATG9B mutation does not increase anxiety-like behaviors.

To assess short- and long-term memory in WT and knockin mice, a novel object recognition test was performed. No significant differences were observed between WT and knockin mice in either short-term (p=0.437, t=0.79) or long-term memory (p=0.650, t=0.460). Similarly, the novel location recognition test, which also assesses short- and long-term memory, showed no significant differences (p=0.710, t=0.376; p=0.328, t=-0.996) between the two groups (Figure S3B and C). These results indicate that ATG9B truncation does not cause short- or long-term memory defects contributing to intellectual disability observed in patients.

The socialization and social preference indices were also similar between the experimental groups (p=0.336, t=-0.982), suggesting that the ATG9B mutation does not affect social behavior (supplementary figure 2-D The social memory test further confirmed the absence of significant differences between WT and knock-in mice (p=0.701, t=0.391).

The marble burying test did not reveal significant differences (p=0.203, t=1.304) in the number of buried marbles between WT and knock-in mice, indicating that the ATG9B truncation does not elicit stereotypical behaviors (Figure S2B).

In the passive avoidance test, Kaplan-Meier survival analysis revealed a trend towards significance between WT and ATG9B-KI mice (Log Rank (Mantel-Cox) Chi-Square=3.072, p=0.08) (Figure 8B), with the KI group displaying a higher tendency to enter the dark compartment compared to the WT group. This suggests a deficit in fear memory recall, as ATG9B-KI mice failed to recollect the association between the dark chamber and the previously encountered aversive stimulus.

In conclusion, the ATG9B mutation does not influence anxiety, repetitive behaviors, or social behaviors, nor does it affect explicit memory functions dependent on the hippocampus or perirhinal cortex. However, the ATG9B mutation may impair amygdala-dependent fear memory.

## Discussion

In this study, we define a novel pathogenic mutation in the ATG9B gene first identified in pediatric patients with neurodevelopmental disorders. Due to the deletion, the C-terminal cytosolic domain of ATG9B protein is deleted, resulting in a truncated ATG9B. We suggest this is a loss of function mutation, as truncated ATG9B protein is unstable and degraded when expressed in cells.

Human ATG9B expression is notably high in the placenta, an organ crucial for maternal and fetal health. Proper placental development is essential for fetal growth, and placental abnormalities are often linked to neurodevelopmental disorders in children (Baschat 2011; Rosenfeld 2021). Our data indicate that ATG9B expression increases after cytotrophoblasts fuse to form syncytiotrophoblasts. BeWo cells, a model for syncytiotrophoblasts in cell culture (Orendi et al. 2010) showed increased ATG9B expression upon induction of syncytialization.

ATG9B TR is localized to perinuclear vesicles as large puncta, while failing to localize in the cytoplasmic vesicles at the periphery like ATG9B WT. Additionally, the expression of ATG9B TR was very low compared to WT which was observed in all transfections we performed. To assess the stability of truncated ATG9B, both WT and truncated ATG9B are ectopically expressed in human cell lines. The protein level of the truncated form was significantly lower than that of the WT in both western blot and immunofluorescence assays while their RNA levels were similar suggesting that the truncated protein is unstable in cells. Recent structural data indicate that the C-terminal cytosolic part of ATG9B is essential for its 3D conformation and homo-trimerization, which is critical for its lipid scramblase function. Although we did not directly study the effect of truncation on ATG9B structure and function, the loss of the C-terminal likely renders the protein non-functional.

The presented ATG9B mutation in this work is not linked to any Mendelian disease. Considering the instability of the truncated protein *in vitro*, and the need for *in-vivo* characterization we modeled the mutation in mice. For phenotypic characterization, we addressed the effect of the mutation on viability, fertility, and the structural and transcriptional landscape of the placenta. Histomorphometry analysis revealed no significant structural differences between heterozygous and homozygous placentas in major structures (decidua, junctional, and labyrinth zones). Despite these findings, more detailed phenotyping focusing on syncytiotrophoblast cells may be required. Immunohistochemistry on human term placenta showed that ATG9B is mainly expressed in syncytiotrophoblasts, marking the first report of ATG9B protein expression in the placenta.

Given the elevated expression of ATG9B in syncytiotrophoblasts, we investigated potential variations in its expression at the transcript level during development and compared WT and mutant mice placentas. Genes involved in syncytiotrophoblast differentiation were evaluated in parallel. We found no significant difference in Atg9b expression between control or mutant placenta samples, yet Atg9b expression was higher at 18.5 dpc compared to 15.5 dpc. The expression of syncytiotrophoblast genes did not show an alteration between placentas of different genotypes. However, here we provide evidence of their varied expression in 18.5 and 15.5 dpc. We concluded that the truncation of Atg9b in a mouse model did not disrupt the gross placenta structure and did not affect the gene expression during syncytiotrophoblast formation.

To address the neurobehavioral aspects of the disorder, we evaluated the knock-in mice in a series of behavioral tests. Despite testing multiple behavioral phenotypes such as learning, memory, and anxiety, we did not observe significant differences. Fear memory, an amygdala-dependent response involving complex brain regions showed a reduction trend in knock-in mice.

Overall, we propose that the ATG9B mutation described in this study is pathogenic in humans and causes a neurodevelopmental disorder with Mendelian inheritance. The truncated ATG9B protein is unstable when expressed in mammalian cells. However, the knock-in mice did not reveal a significant phenotype other than the reduced fear memory trend. There are several potential explanations for the lack of phenotype. Although mouse models are excellent resources for modeling human mutations, there are important differences between human and mouse brains (Hodge et al. 2019; M. H. Kim et al. 2023; Wong et al. 2023). It may be difficult to generate or detect phenotypes similar to intellectual disability observed in patients. The mouse models of human neurodevelopmental diseases do not always reflect on the mouse and result in mild or no phenotype (Ehlers et al. 2023) Moreover, while mouse placenta is frequently used in studies due to shared features, mouse placenta has substantial differences with human placenta in both histological and transcriptomic aspects (Hemberger, Hanna, and Dean 2019; Soncin et al. 2018). Even though the expression of the genes that function in syncytiotrophoblast formation did not change, there may be subtle phenotypical changes in mutant mice that we could now detect with the methods employed in this work. To address this thoroughly a transcriptome and proteome analysis could be performed. Lastly, we cannot exclude the possibility that ubiquitous expression of Atg9a in all tissues including the placenta could compensate for the loss or decrease of Atg9b function in Atg9B knockin mice.

## Supporting information

Kilic et al. supplementary

## ASSOCIATED CONTENT

### Supporting Information

Supporting information including tables and figures mentioned in the text was provided

### Author contributions

The manuscript was written through contributions of all authors. All authors have given approval to the final version of the manuscript.

## NOTES

The authors declare no competing financial interest.

## ACKNOWLEDGEMENTS

We thank Liubovi Sopco and Nazlıcan Kesmik for technical assistance.

## References

1. Aoki, Aiko et al. 2018. “Trophoblast-Specific Conditional Atg7 Knockout Mice Develop Gestational Hypertension.” American Journal of Pathology 188(11): 2474–86. http://ajp.amjpathol.org/article/S0002944018303663/fulltext (November 14, 2022).

2. Ashkenazy, Haim et al. 2016. “ConSurf 2016: An Improved Methodology to Estimate and Visualize Evolutionary Conservation in Macromolecules.” Nucleic Acids Research 44. https://academic.oup.com/nar/article-abstract/44/W1/W344/2499373 (August 26, 2024).

3. Bailey, Craig H., Eric R. Kandel, and Kristen M. Harris. 2015. “Structural Components of Synaptic Plasticity and Memory Consolidation.” Cold Spring Harbor Perspectives in Biology 7(7): 1–29. /pmc/articles/PMC4484970/ (August 13, 2024).

4. Baschat, A. A. 2011. “Neurodevelopment Following Fetal Growth Restriction and Its Relationship with Antepartum Parameters of Placental Dysfunction.” Ultrasound in obstetrics & gynecology : the official journal of the International Society of Ultrasound in Obstetrics and Gynecology 37(5): 501–14. https://pubmed.ncbi.nlm.nih.gov/21520312/ (October 17, 2023).

5. Brown, Richard E., Lianne Stanford, and Heather M. Schellinck. 2000. “Developing Standardized Behavioral Tests for Knockout and Mutant Mice.” ILAR journal 41(3): 163–74. https://pubmed.ncbi.nlm.nih.gov/11406708/ (August 13, 2024).

6. Chen, Siwei et al. 2023. “A Genomic Mutational Constraint Map Using Variation in 76,156 Human Genomes.” Nature 2023 625:7993 625(7993): 92–100. https://www.nature.com/articles/s41586-023-06045-0 (August 12, 2024).

7. Cheng, Yan, Xingcong Ren, William N. Hait, and Jin Ming Yang. 2013. “Therapeutic Targeting of Autophagy in Disease: Biology and Pharmacology.” Pharmacological Reviews 65(4): 1162. /pmc/articles/PMC3799234/ (July 26, 2024).

8. Chiduza, George N. et al. 2023. “ATG9B Is a Tissue-Specific Homotrimeric Lipid Scramblase That Can Compensate for ATG9A.” Autophagy. https://www.tandfonline.com/doi/full/10.1080/15548627.2023.2275905 (November 14, 2023).

9. Claude-Taupin, Aurore et al. 2018. “ATG9A Is Overexpressed in Triple Negative Breast Cancer and Its In Vitro Extinction Leads to the Inhibition of Pro-Cancer Phenotypes.” Cells 7(12). https://pubmed.ncbi.nlm.nih.gov/30563263/ (August 8, 2024).

10. Collier, Jack J. et al. 2021. “Developmental Consequences of Defective ATG7-Mediated Autophagy in Humans.” New England Journal of Medicine 384(25): 2406–17. https://www.nejm.org/doi/full/10.1056/NEJMoa1915722 (July 7, 2024).

11. Ehlers, Julia Sophie et al. 2023. “Morphological and Behavioral Analysis of Slc35f1-Deficient Mice Revealed No Neurodevelopmental Phenotype.” Brain Structure and Function 228(3–4): 895– 906. https://link.springer.com/article/10.1007/s00429-023-02629-8 (October 15, 2024).

12. Elmore, Susan A. et al. 2022. “Histology Atlas of the Developing Mouse Placenta.” Toxicologic Pathology 50(1): 60–117.

13. Eltokhi, Ahmed, Barbara Kurpiers, and Claudia Pitzer. 2020. “Behavioral Tests Assessing Neuropsychiatric Phenotypes in Adolescent Mice Reveal Strain- and Sex-Specific Effects.” Scientific Reports 2020 10:1 10(1): 1–15. https://www.nature.com/articles/s41598-020-67758-0 (August 13, 2024).

14. Erguven, Mehmet, Seval Kilic, Ezgi Karaca, and M. Kasim Diril. 2023. “Genetic Complementation Screening and Molecular Docking Give New Insight on Phosphorylation-Dependent Mastl Kinase Activation.” Journal of Biomolecular Structure and Dynamics 41(17): 8241–53.

15. Guardia, Carlos M. et al. 2020. “Structure of Human ATG9A, the Only Transmembrane Protein of the Core Autophagy Machinery.” Cell Reports 31(13): 107837. http://www.cell.com/article/S2211124720308184/fulltext (November 13, 2023).

16. Hemberger, Myriam, Courtney W. Hanna, and Wendy Dean. 2019. “Mechanisms of Early Placental Development in Mouse and Humans.” Nature Reviews Genetics 2019 21:1 21(1): 27–43. https://www.nature.com/articles/s41576-019-0169-4 (October 15, 2024).

17. Hiz, Semra et al. 2022. “VARS1 Mutations Associated with Neurodevelopmental Disorder Are Located on a Short Amino Acid Stretch of the Anticodon-Binding Domain.” Turkish Journal of Biology 46(6): 458–64.

18. Hodge, Rebecca D. et al. 2019. “Conserved Cell Types with Divergent Features in Human versus Mouse Cortex.” Nature 2019 573:7772 573(7772): 61–68. https://www.nature.com/articles/s41586-019-1506-7 (September 4, 2024).

19. Hung, Tai Ho et al. 2013. “Autophagy in the Human Placenta throughout Gestation.” PLoS ONE 8(12): 1–11.

20. Kandel, Eric R., Yadin Dudai, and Mark R. Mayford. 2014. “The Molecular and Systems Biology of Memory.” Cell 157(1): 163–86.

21. Kang, Mi Ran et al. 2009. “Frameshift Mutations of Autophagy-related Genes ATG2B, ATG5, ATG9B and ATG12 in Gastric and Colorectal Cancers with Microsatellite Instability.” The Journal of Pathology 217(5): 702–6. https://onlinelibrary.wiley.com/doi/full/10.1002/path.2509 (July 7, 2024).

22. Kim, Mean Hwan et al. 2023. “Target Cell-Specific Synaptic Dynamics of Excitatory to Inhibitory Neuron Connections in Supragranular Layers of Human Neocortex.” eLife 12. https://pubmed.ncbi.nlm.nih.gov/37249212/ (September 4, 2024).

23. Kim, Myungjin et al. 2016. “Mutation in ATG5 Reduces Autophagy and Leads to Ataxia with Developmental Delay.” eLife 5(JANUARY2016).

24. Klionsky, Daniel J et al. “For Monitoring Autophagy (4Th Edition).”

25. Kuma, Akiko, Masaaki Komatsu, and Noboru Mizushima. 2017. “Autophagy-Monitoring and Autophagy-Deficient Mice.” Autophagy 13(10): 1619–28.

26. Landrum, Melissa J. et al. 2018. “ClinVar: Improving Access to Variant Interpretations and Supporting Evidence.” Nucleic Acids Research 46(D1): D1062–67. 10.1093/nar/gkx1153 (August 23, 2023).

27. Li, Xia, Zhuo Hang Li, Ying Xiong Wang, and Tai Hang Liu. 2023. “A Comprehensive Review of Human Trophoblast Fusion Models: Recent Developments and Challenges.” Cell Death Discovery 2023 9:1 9(1): 1–14. https://www.nature.com/articles/s41420-023-01670-0 (August 28, 2024).

28. Lu, Jinhua et al. 2013. “A Positive Feedback Loop Involving Gcm1 and Fzd5 Directs Chorionic Branching Morphogenesis in the Placenta.” PLoS Biology 11(4).

29. Matoba, Kazuaki et al. 2020. “Atg9 Is a Lipid Scramblase That Mediates Autophagosomal Membrane Expansion.” Nature Structural & Molecular Biology 2020 27:12 27(12): 1185–93. https://www.nature.com/articles/s41594-020-00518-w (November 12, 2023).

30. Matoba, Kazuaki, and Nobuo N. Noda. 2021. “Structural Catalog of Core Atg Proteins Opens New Era of Autophagy Research.” The Journal of Biochemistry 169(5): 517–25. 10.1093/jb/mvab017 (August 26, 2024).

31. Van Meer, Peter, and Jacob Raber. 2005. “Mouse Behavioural Analysis in Systems Biology.” Biochemical Journal 389(Pt 3): 593. /pmc/articles/PMC1180709/ (September 3, 2024).

32. Mehrabi Pour, Mahsa, Mahboobeh Nasiri, Hajar Kamfiroozie, and Mohammad Javad Zibaeenezhad. 2019. “Association of the ATG9B Gene Polymorphisms with Coronary Artery Disease Susceptibility: A Case-Control Study.” Journal of Cardiovascular and Thoracic Research 11(2): 109–15. https://pubmed.ncbi.nlm.nih.gov/31384404/ (November 12, 2020).

33. Modzelewski, Andrew J. et al. 2018. “Efficient Mouse Genome Engineering by CRISPR-EZ Technology.” Nature Protocols 13(6): 1253–74. https://pubmed.ncbi.nlm.nih.gov/29748649/ (April 28, 2021).

34. Noda, Takeshi, Kuninori Suzuki, and Yoshinori Ohsumi. 2002. “Yeast Autophagosomes: De Novo Formation of a Membrane Structure.” Trends in Cell Biology 12(5): 231–35. https://pubmed.ncbi.nlm.nih.gov/12062171/ (October 2, 2024).

35. Nunes, Joao et al. 2016. “ATG9A Loss Confers Resistance to Trastuzumab via C-Cbl Mediated Her2 Degradation.” Oncotarget 7(19): 27599–612. https://www.oncotarget.com/article/8504/text/ (August 8, 2024).

36. Oral, Ozlem, Ozlem Yedier, Seval Kilic, and Devrim Gozuacik. 2017. “Involvement of Autophagy in T Cell Biology.” Histology and Histopathology 32(1): 11–20. https://pubmed.ncbi.nlm.nih.gov/27225864/ (November 8, 2020).

37. Orendi, Kristina et al. 2010. “The Choriocarcinoma Cell Line BeWo: Syncytial Fusion and Expression of Syncytium-Specific Proteins.” Reproduction 140(5): 759–66. www.reproduction-online.org (April 19, 2021).

38. Rosenfeld, Cheryl S. 2021. “The Placenta-Brain-Axis.” Journal of Neuroscience Research 99(1): 271–83. https://onlinelibrary.wiley.com/doi/full/10.1002/jnr.24603 (September 7, 2023).

39. Saito, S., and A. Nakashima. 2013. “Review: The Role of Autophagy in Extravillous Trophoblast Function under Hypoxia.” Placenta 34(SUPPL): S79–84. 10.1016/j.placenta.2012.11.026.

40. Schindelin, Johannes et al. 2012. “Fiji: An Open-Source Platform for Biological-Image Analysis.” Nature Methods 2012 9:7 9(7): 676–82. https://www.nature.com/articles/nmeth.2019 (October 15, 2024).

41. Shi, Wei et al. 2004. “Choroideremia Gene Product Affects Trophoblast Development and Vascularization in Mouse Extra-Embryonic Tissues.” Developmental Biology 272(1): 53–65. https://pubmed.ncbi.nlm.nih.gov/15242790/ (September 2, 2024).

42. Simmons, David G. et al. 2008. “Early Patterning of the Chorion Leads to the Trilaminar Trophoblast Cell Structure in the Placental Labyrinth.” Development 135(12): 2083–91. 10.1242/dev.020099 (June 10, 2024).

43. Soares, Michael J., Khursheed Iqbal, and Keisuke Kozai. 2017. “HYPOXIA AND PLACENTAL DEVELOPMENT.” Birth defects research 109(17): 1309. /pmc/articles/PMC5743230/ (September 5, 2021).

44. Soncin, Francesca et al. 2018. “Comparative Analysis of Mouse and Human Placentae across Gestation Reveals Species-Specific Regulators of Placental Development.” Development (Cambridge) 145(2).

45. Thul, Peter J., et al. 2017. “A Subcellular Map of the Human Proteome.” Science 356(6340). https://www.science.org/doi/10.1126/science.aal3321 (August 12, 2024).

46. Turco, Margherita Y., and Ashley Moffett. 2019. “Development of the Human Placenta.” Development 146(22).

47. Uhlén, Mathias, et al. 2015. “Tissue-Based Map of the Human Proteome.” Science 347(6220). https://www.science.org/doi/10.1126/science.1260419 (August 12, 2024).

48. Wong, Hovy Ho Wai, Christina You Chien Chou, Alanna Jean Watt, and Per Jesper Sjöström. 2023. “Comparing Mouse and Human Brains.” eLife 12. /pmc/articles/PMC10332809/ (September 4, 2024).

49. Yamada, Takahiro et al. 2005. “Endothelial Nitric-Oxide Synthase Antisense (*NOS3AS*) Gene Encodes an Autophagy-Related Protein (APG9-Like2) Highly Expressed in Trophoblast.” Journal of Biological Chemistry 280(18): 18283–90. http://www.jbc.org/lookup/doi/10.1074/jbc.M413957200 (April 18, 2020).

50. Yamaguchi, Junji et al. 2018. “Atg9a Deficiency Causes Axon-Specific Lesions Including Neuronal Circuit Dysgenesis.” Autophagy 14(5): 764–77. https://www.tandfonline.com/doi/full/10.1080/15548627.2017.1314897 (October 1, 2020).

51. Yang, Ying, and Daniel J. Klionsky. 2020. “Autophagy and Disease: Unanswered Questions.” Cell Death and Differentiation 27(3): 858–71. 10.1038/s41418-019-0480-9.

52. Young, Andrew R.J. et al. 2006. “Starvation and ULK1-Dependent Cycling of Mammalian Atg9 between the TGN and Endosomes.” Journal of Cell Science 119(18): 3888–3900.

