## Supplementary material for "Characterization of a novel neurodevelopmental rare disease caused by a mutation within the autophagy gene *ATG9B*": Kilic et al. supplementary

**Supplementary Figures**

Table 1 The list of oligonucleotides used in this study.

| Primer name | Purpose | 5'>3' sequence |
|---|---|---|
| Primer 1 | Human ATG9B Sanger FOR | CCCATTGGTTCTAAGCGCC |
| Primer 2 | Human ATG9B Sanger REV | GTTAGGGGACGAGGCATGG |
| Primer 3 | pcDNA3.1 FLAG ATG9B FOR primer 1 | AAAGACGATGACGACAAGGTGAGCCGA ATGGGC |
| Primer 4 | pcDNA3.1 FLAG ATG9B FOR primer 2 | GATCAAGCTTATGGACTACAAAGACGAT GACGAC |
| Primer 5 | pcDNA3.1 FLAG ATG9B WT REV primer 1 | GATCCTCGAGTCAGTCAGTGCAAGAGG |
| Primer 6 | pcDNA3.1 FLAG ATG9B TR REV primer 1 | TTGCCGTCCTCCGCACGAGGCCTCAGT CTG |
| Primer 7 | pcDNA3.1 FLAG ATG9B TR REV primer 2 | TGTACGGTGGGAGGTCTATATA |
| Primer 8 | pcDNA3.1 myc his A subcloning | GATCAAGCTTATGGTGAGCCGAATGGG CTGGGG |
| Primer 9 | pBOBI subcloning | GATCGGATCCATGGACTACAAAGACGAT GACGAC |
| Primer 10 | Knock-in Genotyping FOR | GGTGGTGGAAGAGTGGAGTG |
| Primer 11 | Knock-in Genotyping REV | GATGGAGGGTGGCAAGAACA |
| Primer 12 | Mouse Atg9b RT-PCR FOR | ATGTACCCGAAGGACTCCG |
| Primer 13 | Mouse Atg9b RT-PCR REV | TGGTTGGTTGTTGAAGAGAACAT |
| Primer 14 | Mouse Gcm1 FOR | CCCCAGCAAGTTCCATCAGA |
| Primer 15 | Mouse Gcm1 REV | AAGGCTCACCTCCCGGATT |
| Primer 16 | Mouse Syna FOR | AGATACCCCGATGACCACGTC |
| Primer 17 | Mouse Syna REV | TGAGGATCGTCTGGGTGGAG |
| Primer 18 | Mouse Synb FOR | CCACCACCCATACGTTCAAA |
| Primer 19 | Mouse Synb REV | GGTTATAGCAGGTGCCGAAG |
| Primer 20 | Mouse Cebpa FOR | AAAGCCAAGAAGTCGGTGGAC |
| Primer 21 | Mouse Cebpa REV | CTTTATCTCGGCTCTTGCGC |
| sgRNA 1 | Mouse Atg9b knock-in | GGATCCTAATACGACTCACTATAGGCTT TCAGAGGGACAGACAGGTTTTAGAGCTA GAA |
| sgRNA 2 | Mouse Atg9b knock-in | GGATCCTAATACGACTCACTATAGGCTC TGCACGCTGAGAGAGTGGTTTTAGAGC TAGAA |
| HDR template | Mouse Atg9b knock-in | ATACCAGAAGGACTCCCTTCCTTGCAGT GGCTTTCAGAGGGACAGACAGAattCTgA CTCTCTCAGCGTGCAGAGGATGGAAAG ACCGAACTCTCCTTAATGCGGTTCTC |

### Supplementary Figure 1

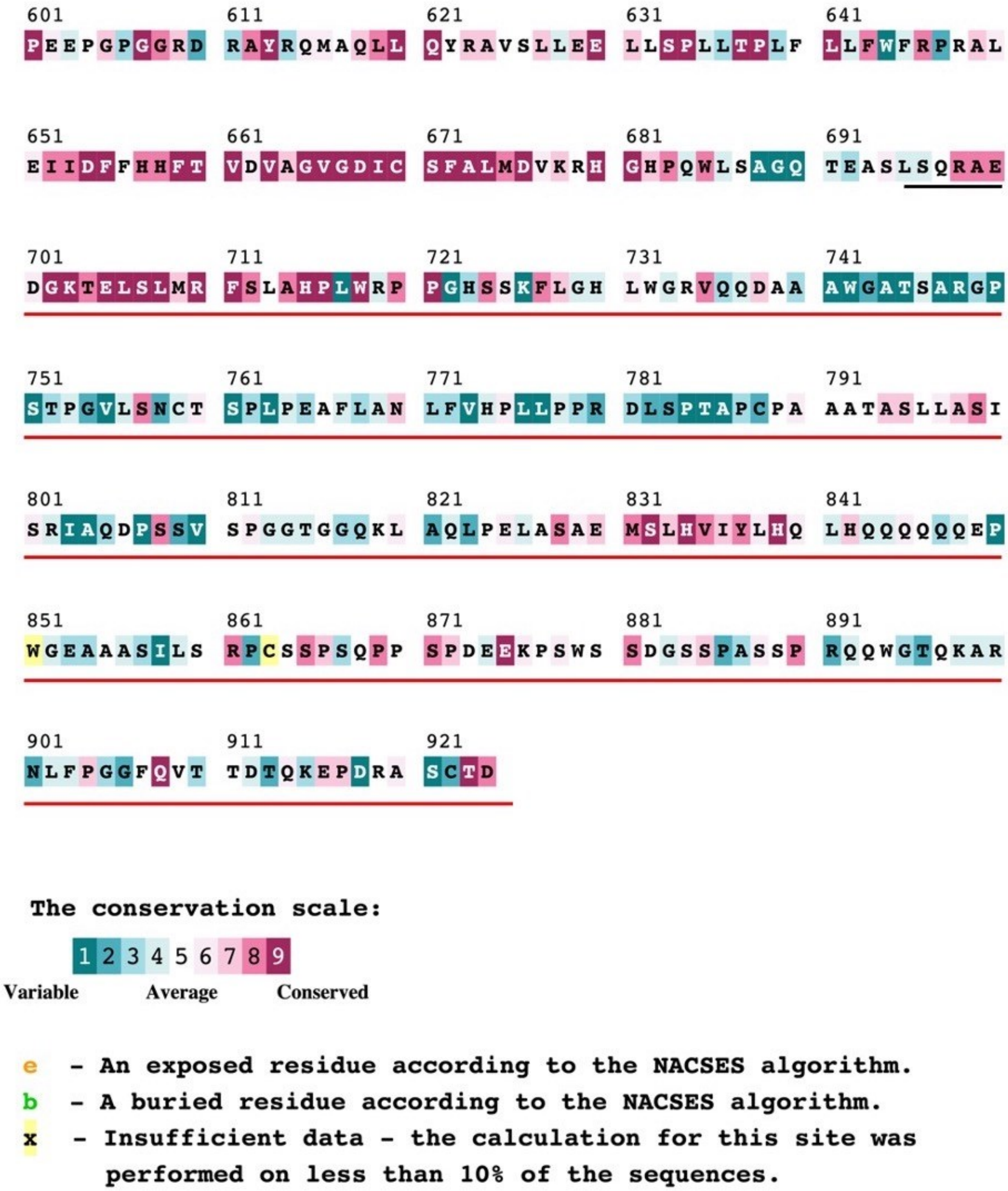


**Supplementary Figure 1** Conservation of C-terminal amino acids of ATG9B, affected by the mutation according to ConSurf. The black line indicates altered amino acids, and the red line indicates deleted amino acids due to the mutation.

### Supplementary Figure 2

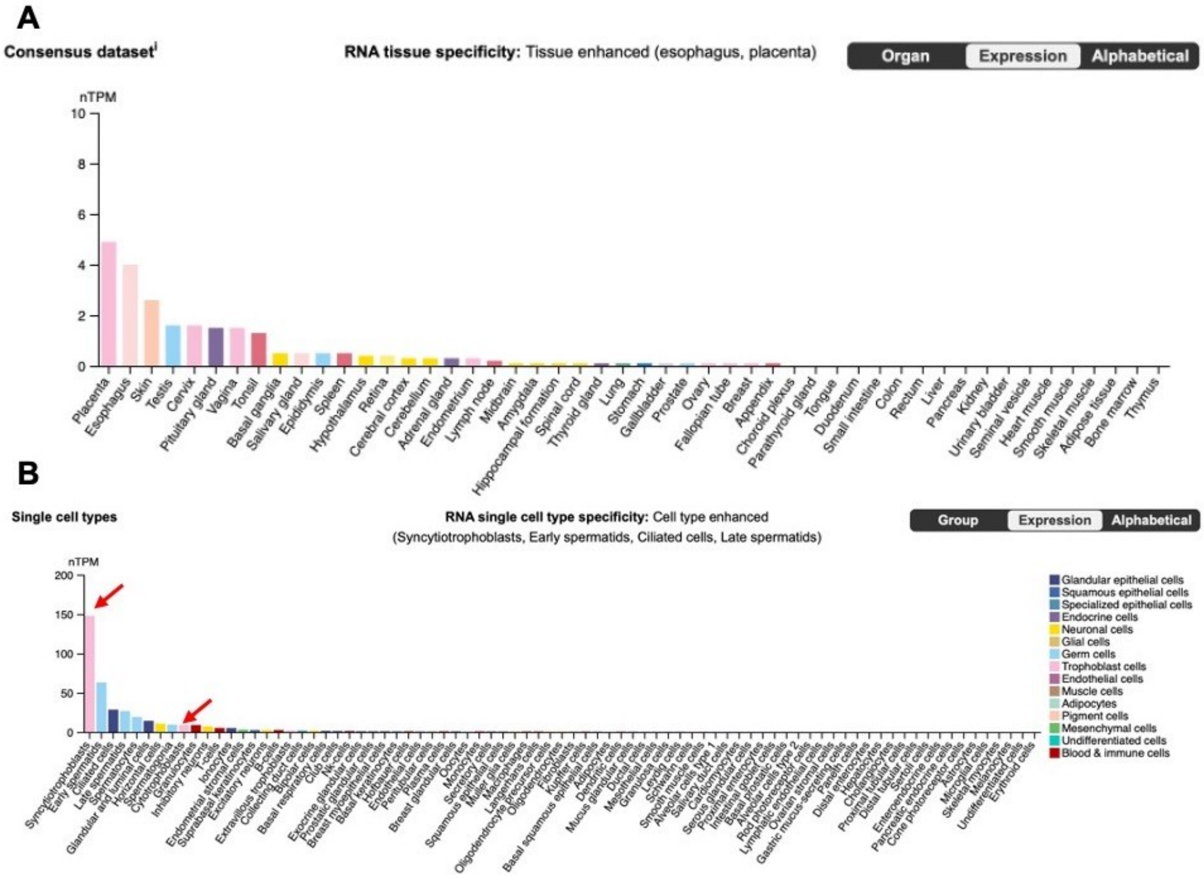


**Supplementary Figure 2** RNA level ATG9B expression among tissues and cell types. A) Among human tissues, the placenta expresses ATG9B the highest. B) Among human cell types, placenta syncytiotrophoblast cells express ATG9B the highest, and cytotrophoblast cells express moderately (Protein Atlas).

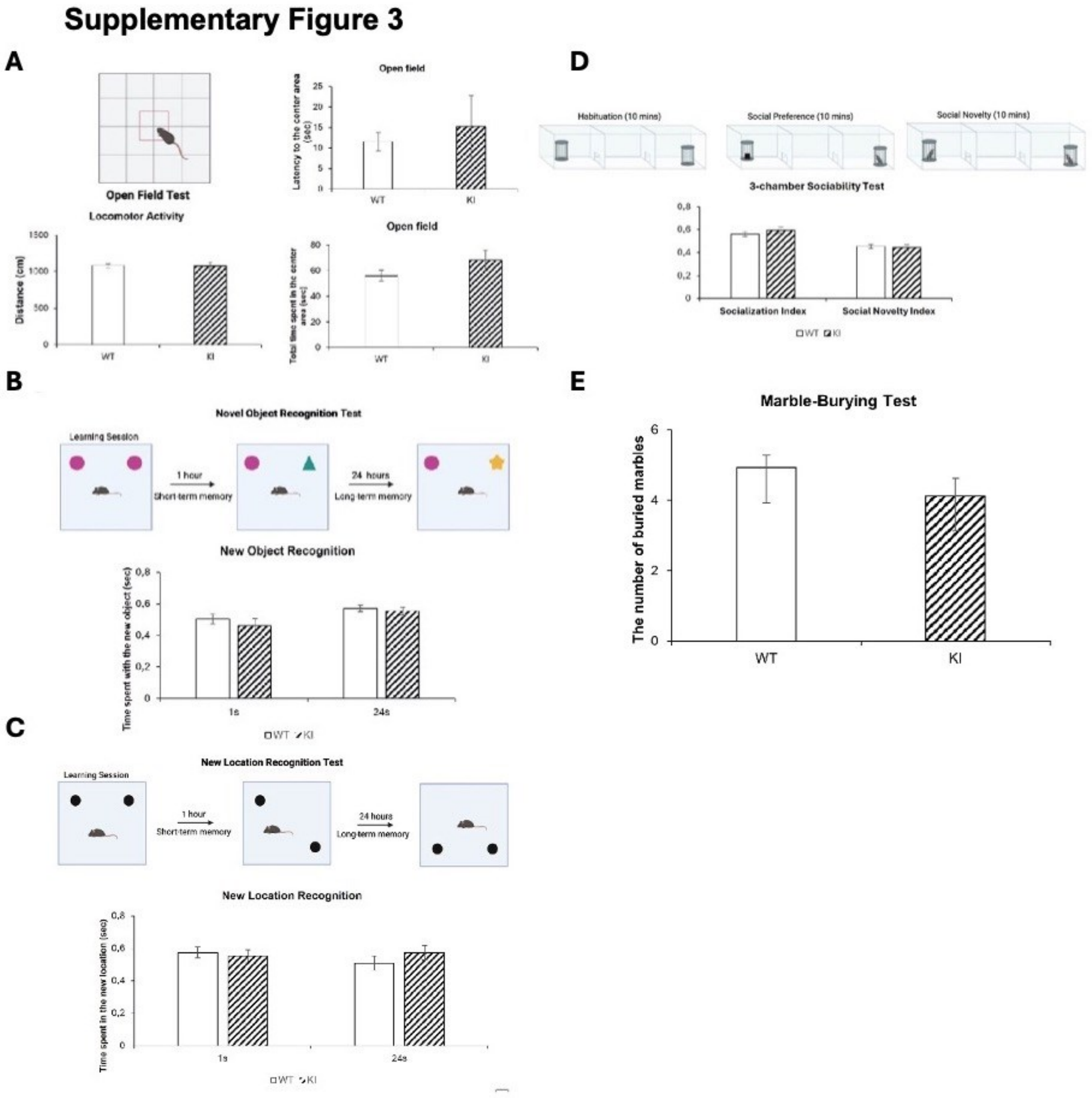


**Supplementary Figure 3** A) Mice were placed in a wooden box (22.5x22.5x30 cm) with a white plexiglass bottom and their behaviors were recorded for 10 minutes. Anxiety-like behavior was assessed by the total time spent in the central 5x5 cm² open area of the box, frequency of entrances into the open area, and latency to first enter the open area. B) Novel object recognition test was conducted by placing identical objects were placed on the opposite corners of the 22.5x22.5x30 cm box. After 10 minutes learning period one of the objects was replaced with a different object. The mice were recorded for 10 minutes. Time spent around the new object, relative to time spent around both objects was analyzed as a measure of memory. The test started one hour after the learning period for the short-term memory, 24 hours after for the long-term memory. C) New location recognition test was conducted in the same setup with B. Instead, the location of the object is changed one hour and 24 hours after the learning trial. C) Three-chamber cage was used for the social preference and social memory test. First day, the mice were habituated for 10 minutes to a 3-chambered box, two opposite chambers containing empty cylindrical cages. The second day a live mouse of the same species and gender was placed in one of the cylindrical cages in one chamber, while a toy (a colored wooden block) was placed in the cylindrical cage in the opposite chamber. The mice were placed in the central chamber allowed to freely explore their surroundings for 5 minutes. Then the doors separating the chambers were removed and the movement of the mice between the chambers was recorded for 10 minutes. The time spent interacting with the cylindrical cages on both sides was measured for the social preference test. Socialization index was calculated by the following formula: the time spent around the cylindrical cage containing the stranger mouse/ the time spent around both cylindrical cages. On the third day, social novelty test was carried out to analyze social memory. A familiar mouse (cage mate) and an unfamiliar (stranger) mouse was placed inside the cages in opposite chambers instead. The social memory index was calculated by the following formula: the time spent around the cylindrical cage containing the unfamiliar mouse/ the time spent around both cages. B) Marble burying test was conducted to examine stereotypical behaviors. Clean bedding (3-4 cm) was placed in the cage, and the mice were habituated to the cage for 30 minutes. Thereafter, 15 marbles were placed symmetrically and in a three-by-five arrangement in a grid pattern in the cage. The mice were left to freely roam in the cage for 15 minutes, after which they were returned to their home cages, and the buried marbles were counted.
